# Monitoring the stability of transgene expression in lettuce using the RUBY reporter

**DOI:** 10.1101/2025.08.17.670745

**Authors:** Beth A. Rowan, Megan Reeves, Claire Hays, Cana Shirley, Wathsala Rajapakse, Katrine A. Taran, Tawni Bull, Dylan A. Wong, Richard W. Michelmore

## Abstract

Nearly four decades after the first transgenic lettuce was reported, constructs for stable transgene expression remain limited. Notably, the 35S promoter from the Cauliflower Mosaic Virus (35S), which drives strong expression of transgenes in several plant species, has often shown silencing and instability in lettuce. Other promoter/terminator combinations that are commonly used in plant expression vectors have not been extensively studied in lettuce. In this study, we evaluated three different expression constructs in two different horticultural types of lettuce using the non-invasive RUBY reporter, which allowed for the monitoring of transgene expression throughout the process of regeneration during tissue culture, throughout development of the primary transgenics, and in two subsequent sexual generations. The LsUBI promoter/terminator combination resulted in strong, uniform expression throughout regeneration, during growth of the primary transgenics, and in both subsequent generations. The AtUBI promoter/tRBCS combination showed slightly lower levels of expression and intermediate levels of silencing, while the 35S promoter/tHSP combination showed both initial strong expression and frequent silencing. Therefore, our data show that the LsUBI promoter/terminator combination provides strong, uniform expression that is unlikely to result in silencing and that the AtUBI promoter/tRBCS combination is an additional option for stable expression of transgenes in lettuce, especially if an intermediate expression level is desired.

## Introduction

The ability to introduce genes of interest has a wide variety of potential applications in lettuce, including genome editing (Bertier et al., 2018; Cardi et al., 2017; Kwon et al., 2023; X. Li et al., 2021), engineering traits of importance to growers (Chen et al., 2017; Park et al., 2005; Shelake et al., 2022) and consumers (Dias & Ortiz, 2012; Lim et al., 2008; Yabuta et al., 2013), and for the production of biopharmaceuticals (Boyhan & Daniell, 2011; Fischer et al., 2004; Mohebodini et al., 2014) and other compounds (Hirai et al., 2011). Post-transcriptional gene silencing (PTGS), which likely evolved to defend plants against viruses (Kasschau & Carrington, 1998; H. W. Li et al., 1999; Vance & Vaucheret, 2001), is often triggered in response to transgenes (Béclin et al., 2002; Vaucheret et al., 1998), turning off their expression and obstructing their intended purpose. Long-term silencing can then be maintained through methylation of transgenes and/or their promoter sequences.

Previous studies of transgenic lettuce indicated that the promoter used for transgene expression as a key factor affecting the likelihood of silencing. McCabe et al. (1999) found that only 34% of T_1_ lettuce transgenics with the glufosinate ammonium herbicide resistance gene, *bar*, expressed from the 35S promoter were resistant to the herbicide. The herbicide resistance declined further until only 11% of the 35S-*bar* plants were resistant in the T_3_ generation, while the percentage of herbicide resistant plants remained at 95-100% for three generations when the *bar* gene was expressed from a truncated promoter from the plastocyanin (*petE*) gene from pea. Several previous studies also reported either high levels of silencing or reduced/nonexistent levels of expression for transgenes expressed from the 35S promoter in lettuce (Dinant et al., 1997; Dubois et al., 2005; Hirai et al., 2011; Niki et al., 2001; Okumura et al., 2016; Pang et al., 1996). Okumura et al. (2016) demonstrated that silencing of sGFP in lettuce was associated with DNA methylation at 54-92% of the cytosines in the 35S promoter. Hirai *et al*. (2011) found that none of three lines propagated to the T_2_ generation had detectable expression of a miraculin transgene expressed from the 35S promoter, while all three lines had detectable miraculin when the LsUBI promoter was used (Hirai et al., 2011). In some cases, silencing has been overcome through other molecular processes. For example, Niki et al. (2001) initially found no change in the plant phenotype when a GA oxidase transgene with a 35S promoter paired with the tNOS terminator was used, but did observe the desired altered phenotype when a translational enhancer from tobacco mosaic virus was added. Other studies using the 35S promoter to drive transgene expression in lettuce did not report silencing, or only minimal silencing (de Toledo Thomazella et al., 2016; Goto et al., 2000; Park et al., 2005; Torres et al., 1993), but either did not report how many of the initial T_1_ plants that were generated did not show expression or did not report expression beyond the T_2_ generation.

The optimal promoter/terminator design for transgene expression remains a high priority for engineering desirable traits in lettuce. However, it is difficult to compare and evaluate expression strategies from the literature because most previous studies used only one or rarely two promoters, only one cultivar or horticultural type, and varied in the number of generations over which transgene expression was tracked. Also, silencing has been observed to be variable across development (Pang et al., 1996), and therefore differences in the timing of measurement of gene expression (whether through mRNA, protein, or phenotyping) may affect the determination of whether a particular transgene expression construct is effective.

In this study, we employed the RUBY reporter (He et al., 2020) to monitor expression over three generations using three different transgene constructs in two domesticated lettuce cultivars representing different horticultural types (Figure 1). We tested the 35S promoter and the LsUBI promoter because previous work reported that the 35S promoter was prone to instability, while the lettuce polyubiquitin (LsUBI) promoter was more stable (Hirai et al., 2011). We also tested the *Arabidopsis thaliana* polyubiquitin 10 (AtUBI10) promoter because although it is commonly used in plant expression constructs, there are limited data on its stability in lettuce. The AtUBI10 promoter has been used successfully to drive the expression of Cas9 for genome editing in lettuce (Riu et al., 2023) and to express the growth regulator GRF5 (Pan et al., 2022), but neither of these studies specifically examined the expression of the transgenes either within the primary transgenics or over subsequent generations. Unlike previously developed reporter genes, the visual readout of the RUBY marker requires no specialized equipment, chemical treatments, or destructive sampling. Gene expression can be monitored by simple visual scoring for the presence of brightly colored betalain pigments. Thus, transgene silencing can be monitored throughout the tissue culture regeneration process, during development within the lifetime of individual plants, and across multiple sexual generations.

**Figure 1.**
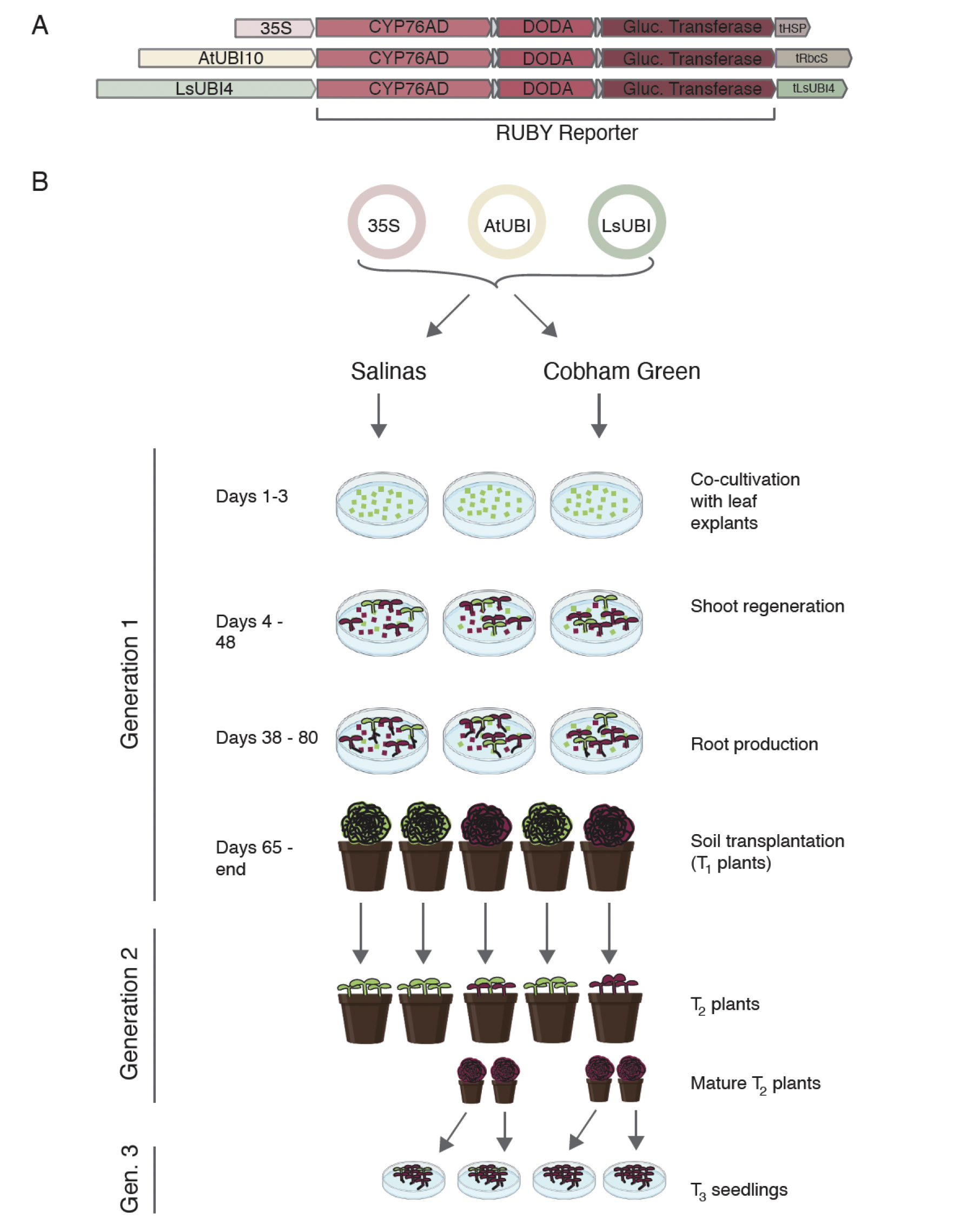
Construct and Experimental Design. A) Schematic illustrations showing the transcriptional units for the three different RUBY constructs. B) The three different RUBY expression constructs shown in (A) were transformed into the domesticated lettuce (*Lactuca sativa*) cultivars ‘Salinas’ and ‘Cobham Green’ and monitored for RUBY expression during tissue culture regeneration, subsequent growth of the T_2_ and T_3_ generation individuals.

## Materials and Methods

### Vectors and Plasmid Construction

The 35S::RUBY-tHSP construct (JD849) was provided by Juan Debernardi (UC Davis Plant Transformation Facility), who obtained the 35S::RUBY-tHSP plasmid from Addgene (https://www.addgene.org/160908/) and modified it to change the plant selectable marker from hygromycin to kanamycin using In-Fusion cloning (Takara Biosciences). The pLsUBI-RUBY-tLsUBI and pAtUBI-RUBY-tRBCS constructs were generated via modular Golden Gate cloning (Weber et al., 2011) as described below.

The single DNA sequence encoding the three proteins of the RUBY cassette was cloned after PCR amplification of the sequence from the 35S::RUBY-tHSP plasmid from Addgene (https://www.addgene.org/160908/) into a Level 0 “SC” vector. The promoter and terminator from the *L. sativa* polyubiquitin 4 gene (LOC111919935) had been previously cloned as Level 0 “PU” and “T” modules, respectively (Bull et al., 2023). The Level 0 “PU” module containing the AtUBI10 promoter, the Level 0 “T” module containing the pea RBCS terminator, Level 2 end linkers and cloning vectors for all Golden Gate levels were obtained from Sylvestre Marillonet (Weber et al., 2011). Golden Gate Level 1 reactions were used to assemble Level 1 modules for LsUBI-RUBY-tLsUBI and AtUBI-RUBY-tRBCS using the Level 0 modules described above and the *BsaI* enzyme. Level 2 binary constructs were generated via Level 2 Golden Gate reactions using the Level 1 pNOS-Nptii-tNOS kanamycin module in the first position adjacent to the left border of the T-DNA, the Level 1 RUBY module in the second position, and the appropriate end-linker (p-ELE-02) into the Level 2 destination vector using the *BpiI/BbsI* enzyme.

### Preparation of bacterial cultures for plant transformation

All plasmids were transformed into *Agrobacterium tumefaciens* strain LBA4404. Initial bacterial cultures were prepared by inoculating 20 mL of MGL medium (5 g/L tryptone, 2.5 g/L yeast extract, 5 g/L NaCl, 5 g/L mannitol, 0.1 g/L MgSO4, 0.25 g/L K2HPO4, 1.2 g/L glutamic acid, 15 g/L sucrose; pH 7.2) supplemented with rifampicin (50 mg/L), kanamycin (50 mg/L), and streptomycin (50 mg/L) with single-colony-derived cultures of *A. tumefaciens* bearing the plasmid of interest. Cultures were incubated overnight in an orbital shaker at 28°C at 180-200 rpm. The following day, subcultures were prepared by inoculating 15 mL of TY medium (5 g/L tryptone, 3 g/L yeast extract; pH 7.2) supplemented with rifampicin (50 mg/L), kanamycin (50 mg/L), and acetosyringone (40 mg/L) with 5 mL of the MGL overnight culture. Cultures were again incubated overnight in an orbital shaker at 28°C at 180-200 rpm. The following morning, cultures were diluted to an OD600 between 0.1 and 0.2. Acetosyringone was added to the final diluted cultures prior to plant transformation at a final concentration of 200 μM.

### Transformation and in vitro tissue culture regeneration of Lactuca sativa cv. Cobham Green and Salinas

Lettuce transformation was carried out using a modification of the method described by Michelmore et al. (1987). Seeds of *L. sativa* cvs. Cobham Green and Salinas were surface sterilized with 20% bleach for 20 min with constant agitation at 250 rpm. Surface-sterilized seeds were rinsed five times with 50 mL of sterile distilled water and sown on 1/2X Hoagland’s medium (0.815 g/L Hoagland modified basal salt mixture [PhytoTech Labs Product ID# H353], 8 g/L PhytoAgar™ [PlantMedia SKU# 40100072-1], pH 5.6–5.8). Seeds were incubated for 4 days in a 24 °C growth room under a 12/12 h light/dark cycle with Honeywell LED lights (Model #SH450505Q2004) providing approximately 8700 lux. After 4 days, explants were prepared by cutting off the apical tip and base of cotyledons while submerged in 20 mL of *A. tumefaciens* suspension culture with OD600 of 0.1 - 0.2. Cotyledon explants were then transferred to SH Co-Cultivation Medium (3.2 g/L Schenk and Hilderbrandt (SH) Basal Salt Mixture, 30 g/L sucrose, 2 mL/L 500X Murashige and Skoog (MS) Vitamins [PhytoTech Labs Product ID# M533], 8 g/L PhytoAgar, pH 5.6-5.8) supplemented with acetosyringone (200 μM), 0.1 mg/L of 6-benzylaminopurine (6-BAP), and 0.1 mg/L of 1-naphthaleneacetic acid (1-NAA) and incubated in the dark for 3 days at 24°C. After 3-4 days on SH Co-Cultivation Medium, the explants were transferred to SH Induction Medium (SHI: 3.2 g/L SH basal salt mixture, 30 g/L sucrose, 2 mL/L 500X MS vitamins, 8 g/L PhytoAgar™, 0.10 mg/L 6-BAP, 0.10 mg/L 1-NAA, 150 mg/L timentin, and 400 mg/L carbenicillin; pH 5.6-5.8). After 14-21 days on SHI, the explants were transferred to fresh SHI and any regenerated shoots were transferred to Shoot Elongation Medium (SHE: 3.2 g/L SH Basal Salt mixture, 30 g/L sucrose, 1X MS vitamins, 8 g/L PhytoAgar™, 0.01 mg/L 6-BAP, 0.05 mg/L 1-NAA, and 150 mg/L Timentin; pH 5.6-5.8). After 28-36 total days post transformation, all regenerating shoots were removed from SHI Medium and any remaining explants and non-regenerating calli were discarded. Regenerating shoots formed rosettes and were kept on SHE Medium until the rosettes were at least 1 cm across and had at least three leaves (typically 24-48 days post transformation), transferring to fresh SHE Medium every 14-21 days. When regenerated shoots met the size selection criteria described above, they were transferred to MS Rooting Medium (MSR: 4.34 g/L Murashige and Skoog [MS] salts, 30 g/L sucrose, 2 mL/L 500X MS vitamins, 8 g/L PhytoAgar™, and 150 mg/L timentin; pH 5.6-5.8), transferring to fresh MSR Medium every 14-21 days until roots at least 1 cm long had formed. When regenerated plantlets exhibited sufficient root development, they were carefully removed from the MSR medium, gently brushing any remaining MSR medium off the roots with a small spatula and transplanted into pots with transparent, closed lids containing autoclaved UC Davis Agronomy Soil mix or Promix mycorhizzae soil. Occasionally, roots had already initiated while the plants were on SHE, in which case they were transplanted to soil without first transferring them to MSR if they were sufficiently developed. Humidity within the pots was gradually reduced by slightly opening the lids of the pots after 3-4 days post-transfer to soil and then removing the lid entirely 7-10 days post-transfer. The plants were then transferred to a greenhouse after an additional 1-2 weeks. Plants were then transplanted to 1-gallon pots 1-2 weeks after transferring to the greenhouse and cultivated under drip irrigation/ fertigation until the plants reached maturity and were ready for harvest. Mesh bags were placed over the inflorescence stalks to prevent seed loss upon shattering and potential cross-fertilization. For the LsUBI-RUBY-tLsUBI Red Cobham Green plants with low self-seed production, crosses were performed by wiping the pollen from wildtype Cobham Green plants on to their stigmas. Individual flower heads that had been crossed were tagged and seeds were collected approximately two weeks after pollination.

### Phenotypic characterization of calli and regenerating shoots

Calli and shoots were evaluated for RUBY expression during the shoot elongation stage. Shoots arising from the same callus were considered conservatively to potentially be from the same transgene integration event. For each transgene integration event, the callus color and the shoot color were recorded. Events were scored as “Green” if no red color was present in the callus or in any one of the regenerating shoots arising from the same callus. Events were scored as “Mixed” if the red color varied within the same callus, or within the same shoot, or among multiple shoots of the same callus. Events were scored as “Red” only if the entire callus and all the shoots arising from it were red.

When plants were scored at the rooting stage, shoots were scored as described above and summary phenotypes were applied to a group of shoots that had originated from the same callus. Roots were scored similarly to shoots, only that the term “White” was assigned as the phenotype if no red color was observed. Some shoots exhibited a uniform phenotype that was not green or red, but a light pink color. This shoot phenotype was designated as “Rosy” and shoots with a Rosy phenotype were scored as Rosy only if all shoots derived from the same transformation event were Rosy. If individual plantlets or multiple plantlets from the same transformation event were mixed between Green and Rosy phenotypes, they were scored as Mixed.

### DNA extraction and PCR genotyping of transgenic T_1_ plants

Leaf samples were obtained from a subset of primary transgenics and DNA was extracted as described in Rowan et al. (2019). PCR genotyping for actin (endogenous gene) and each of two of the cassettes in the T-DNA - the kanamycin resistance cassette and the RUBY cassette - was performed on these samples using the primers described in Table 1. Genotyping results were recorded in a csv file and analyzed in R (R Core Team, 2013), using the UpSetR (Conway et al., 2017) package to summarize the intersection between plants that were positive for the actin gene (control) and each of the two elements of the T-DNA (test).

**Table 1.**
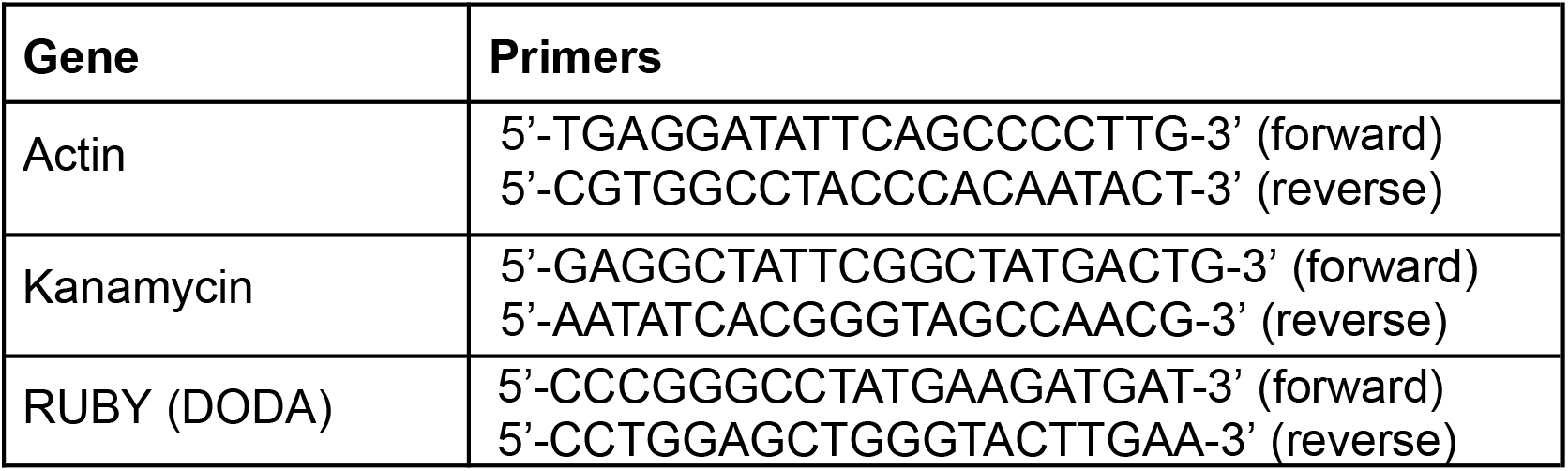
Primers used for PCR screening of primary transgenics:

### Phenotypic analysis of progeny from transgenic plants (T_2_ and T_3_ generations)

For the evaluation of seeds from the primary transgenics (T_1_ generation), eighteen to twenty-one progeny seeds from representative RUBY construct/ parent color categories (Table 2) were sown in 72-pot trays of Promix mycorrhizae soil and kept under a mist system for 20 days. Individual plants were scored as Red, Rosy, Mixed, or Green. Images of the plants were also recorded and the final scores were compiled from both the in-person assessments and the images. The plants were imaged in different lighting conditions, therefore the white balance was adjusted using iPhoto software by selecting the benchtop grating as a neutral grey. Adjustments were performed the same way for all images. A total of 248 individual T_2_ plants representing each construct/phenotype combination were grown to maturity for production of T_3_ seeds. For the evaluation of the T_3_ generation, 20 seeds (or 12-15 if seeds were limited) were first subjected to a brief surface sterilization process (seeds shaken in 10 % bleach with 0.2% Tween-20 for 10 minutes, followed by 4 rinses with deionized water) and then sown on damp Whatman filter paper in Petri plates. After 1 week, the emerged cotyledons were scored phenotypically as Green, Red, Rosy, or Mixed.

**Table 2.** Progeny seeds selected from primary transgenics (T_2_generation).

|  | **Background Genotype** | **T<sub>1</sub> Parent Color<sup>a</sup>** | **Total number of lines** | **Total number of lines with seeds** | **Number of lines selected** |
| --- | --- | --- | --- | --- | --- |
| 35S::RUBY | Cobham Green |  |  |  |  |
|  |  | Green | 8 | 7 | 4 |
|  |  | Mixed | 5 | 5 | 5 |
|  |  | Red | 5 | 2<sup>b</sup> | 0 |
|  | Salinas |  |  |  |  |
|  |  | Green | 20 | 14 | 4 |
|  |  | Mixed | 23 | 19 | 7 |
|  |  | Red | 2 | 2 | 2 |
| AtUBI::RUBY | Cobham Green |  |  |  |  |
|  |  | Green | 14 | 13 | 1 |
|  |  | Mixed | 12 | 10 | 5 |
|  |  | Rosy | 8 | 7 | 4 |
|  |  | Red | 2 | 1<sup>c</sup> | 0 |
|  | Salinas |  |  |  |  |
|  |  | Green | 25 | 12<sup>d</sup> | 4 |
|  |  | Mixed | 16 | 8<sup>e</sup> | 4 |
|  |  | Rosy | 6 | 6 | 6 |
|  |  | Red | 9 | 5<sup>f</sup> | 4 |
| LsUBI::RUBY | Cobham Green |  |  |  |  |
|  |  | Green | 12 | 12 | 4 |
|  |  | Mixed | 5 | 4 | 4 |
|  |  | Red | 6 | 3 | 3 |
|  | Salinas |  |  |  |  |
|  |  | Green | 1 | 1 | 1 |
|  |  | Mixed | 2 | 1 | 1 |
|  |  | Red | 0 | 0 | 0 |

<sup>a</sup> Parent color phenotypes were evaluated at the time of DNA sampling
<sup>b</sup> One line had seed contamination, one line did not have seeds available at the time of sowing for the second generation
<sup>c</sup> One line did not have seeds available at the time of sowing of the second generation
<sup>d</sup> Nine lines died before they could produce seeds
<sup>e</sup> Four lines died before they could produce seeds
<sup>f</sup> Two lines died before they could produce seeds and one line did not have seeds available at the time of sampling

## Results

### Expression of RUBY during early tissue culture regeneration

Red betalain from the RUBY reporter was visible in calli and regenerating shoots for all three expression constructs (35S::RUBY, p35S-RUBY-tHSP; AtUBI::RUBY, pAtUBI1-RUBY-tRBCs; LsUBI::RUBY, pLsUBI-RUBY-tLsUBI) and in each of the two lettuce cultivars (Figure 2). A single callus and its derived shoot(s) were treated as a single transformation event and phenotypes were scored for each transformation event. 35S::RUBY resulted in the lowest percentages of calli with visible RUBY expression (58% of calli for Cobham Green and 54% of calli for Salinas), whereas the percentage of calli with visible RUBY expression was >85% for AtUBI::RUBY and >95% for LsUBI::RUBY in both lettuce backgrounds. LsUBI::RUBY also had the highest percentages of shoots scored as Red (57% in Cobham Green and 42% in Salinas) and the lowest percentages of shoots scored as Green (≤5.2% in both lettuce backgrounds). While there were slight differences between the two lettuce genotypes, they both exhibited a similar trend in expression penetrance (i.e. the percentage of plants scored as Red), with penetrance being the lowest for 35S::RUBY, intermediate for AtUBI::RUBY and highest for LsUBI::RUBY plants.

**Figure 2.**
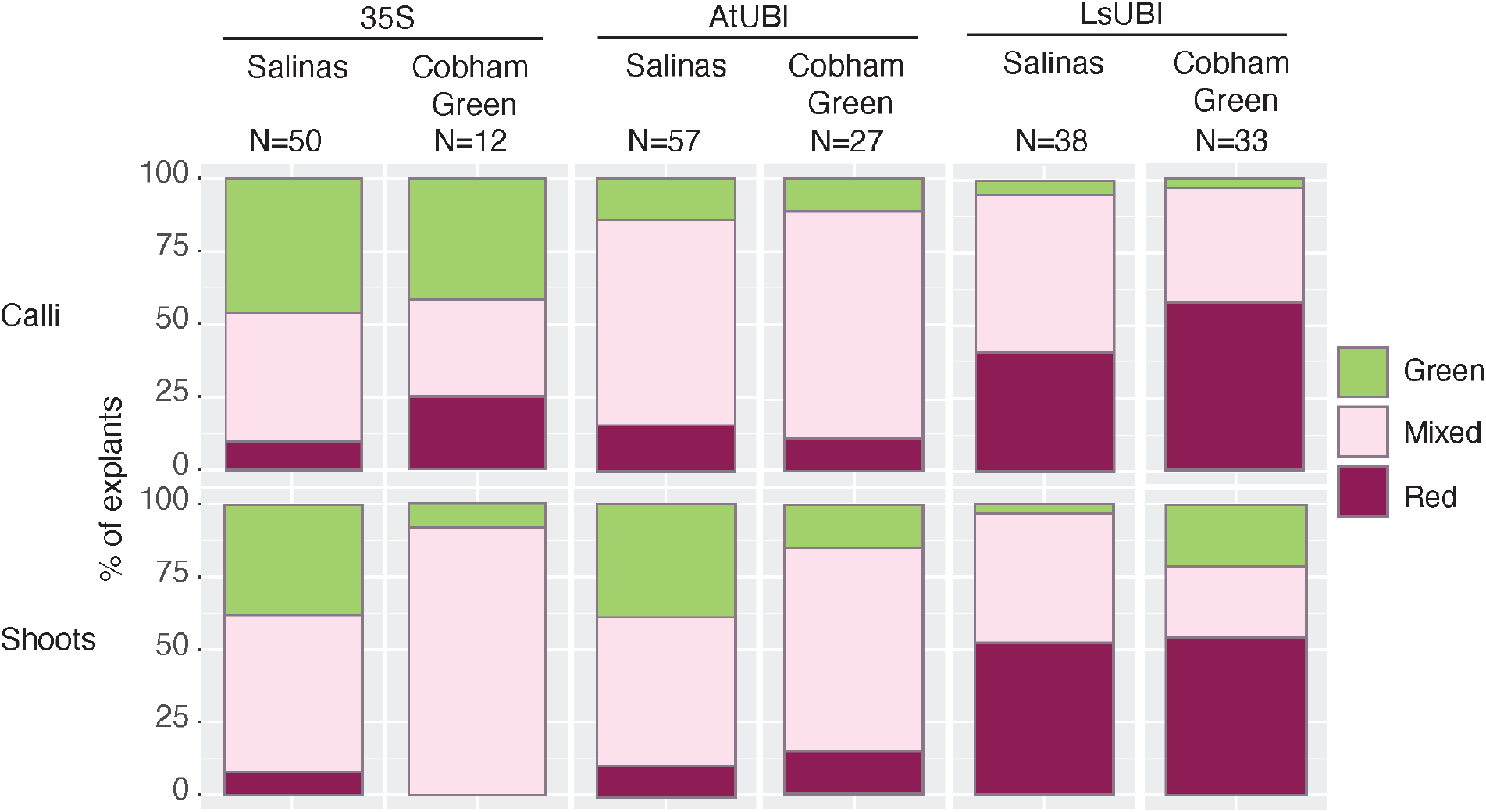
RUBY expression in independent transformation events in calli and regenerating shoots. Calli were scored as “Green” if presenting the typical light yellowish-green color of lettuce calli, “Mixed” for uneven red coloration in the callus, and “Red” only if the callus was uniformly red. Shoots were scored as “Green” if all shoots regenerating from the same callus were green, “Mixed” for either individual shoots with a mixed red/green phenotype or having both red and green shoots regenerating from the same callus, and “Red” if all shoots regenerating from the same callus were red. N=Number of regenerating calli or shoots.

### Expression of RUBY in regenerated T_1_ plants upon transfer to soil and during vegetative growth and reproduction

As regeneration proceeded and roots were initiated and developed, over 50% of the shoots and roots were classified as Red for LsUBI::RUBY for both Cobham Green and Salinas, while less than 40% of the plantlets transformed with AtUBI::RUBY had Red phenotypes in both roots and shoots (Figure 3). At this stage, very few plantlets from the 35S::RUBY transformations exhibited uniform RUBY expression. Out of all the plantlets derived from a total of 45 transformation events for 35S::RUBY in Salinas and Cobham Green, only one showed uniform expression and only in root tissue.

**Figure 3.**
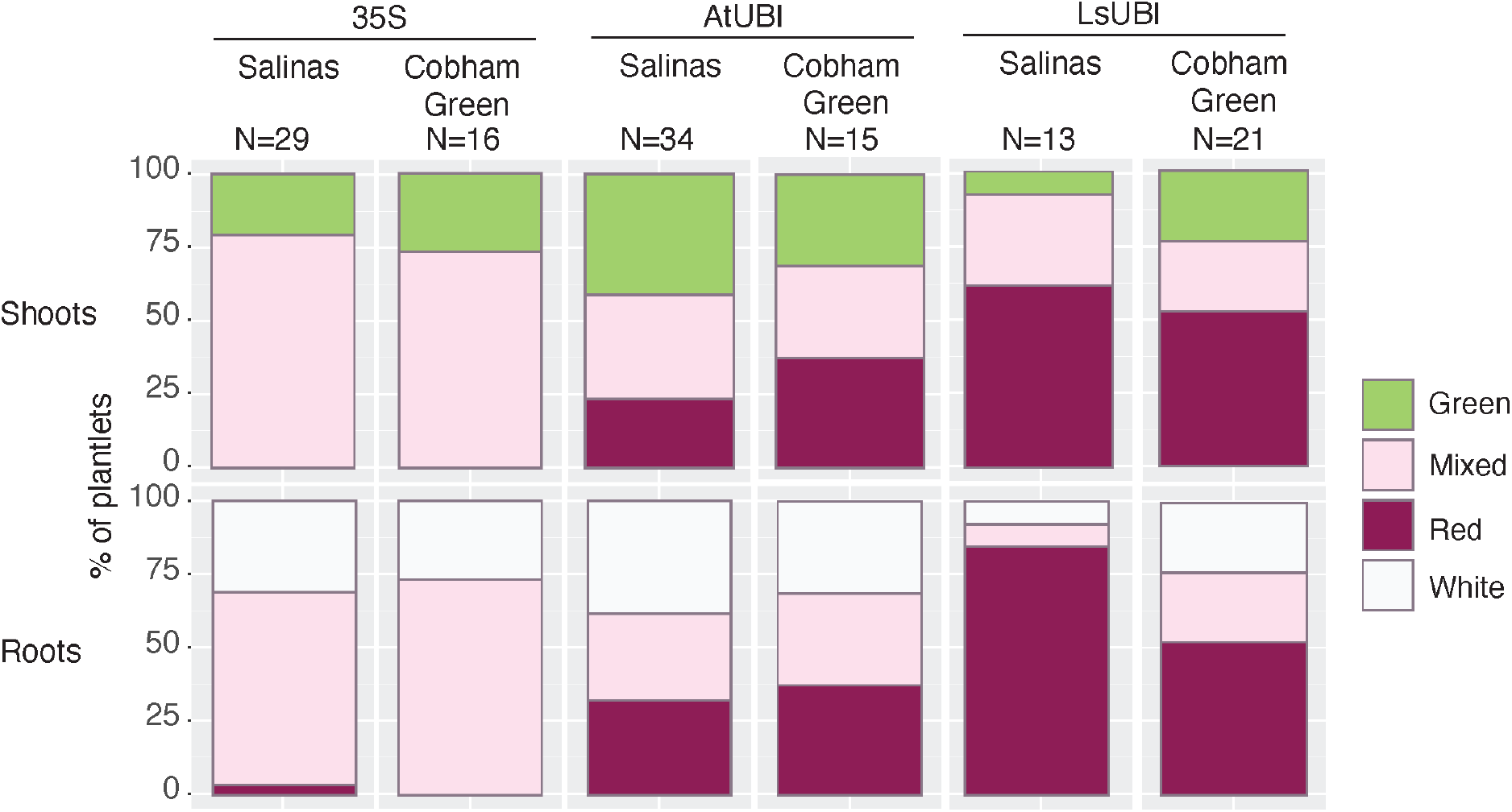
RUBY expression during rooting of regenerated plants from independent transformation events. Shoots were scored as “Green” if all shoots that had originated from the same callus were green, “Mixed” for either individual shoots with a mixed red/green phenotype or both red and green shoots that had regenerated from the same callus, and “Red” if all shoots that had regenerated from the same callus were red. Roots were scored as “White” if the roots of all plants that regenerated from the same callus were white, “Mixed” if the roots had a mix of red and white coloration or if both white and red roots were present among plants that had regenerated from the same callus, and “Red” if all plants that had regenerated from the same callus had uniformly red roots. N=Number of regenerating plants.

Following transplant to soil, LsUBI::RUBY continued to exhibit the most uniform expression of RUBY (Figures 4 and 5). All constructs resulted in the production of at least one individual plant with uniform RUBY expression (Figure 5 and Table 2), even for cases where the expression was inconsistent among shoots derived from the same transformation event and the phenotype for the event was scored as Mixed. However, only two of the individual Red 35S::RUBY Salinas transgenic plants produced seeds and both of the individual Red 35S::RUBY Cobham Green plants were sterile (Table 2). Most 35S::RUBY plants had either a Mixed or Green phenotype.

**Figure 4.**
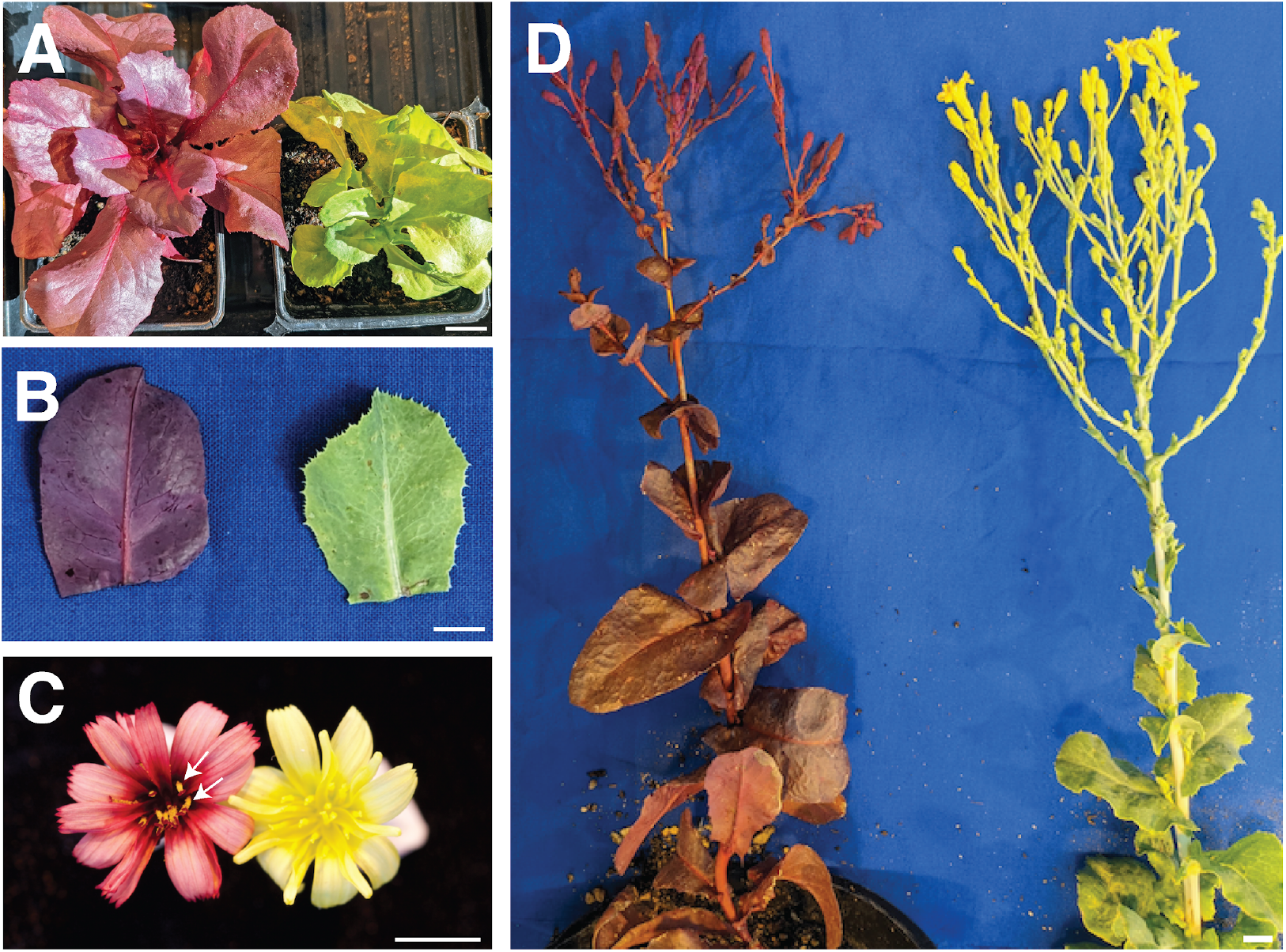
RUBY expression in LsUBI::RUBY *L. sativa* ‘Cobham Green’ T_1_plants. A) Representative plants during early vegetative growth (shown are representative examples of individual LsUBI::RUBY ‘Cobham Green’ plants with either a uniformly red phenotype (left) or entirely green (right) phenotype). B) Representative individual leaves from a flowering stalk with either a uniformly red phenotype (left) or entirely green (right) phenotype after transformation with LsUBI::RUBY. C) Representative individual flowers from LsUBI::RUBY Cobham Green plants with either a uniformly red (left) or entirely green (right) phenotype. White arrows indicate yellow pollen. D) Representative flowering stalks from LsUBI::RUBY Cobham Green plants with either a uniformly red (left) or entirely green (right) phenotype.

**Figure 5.**
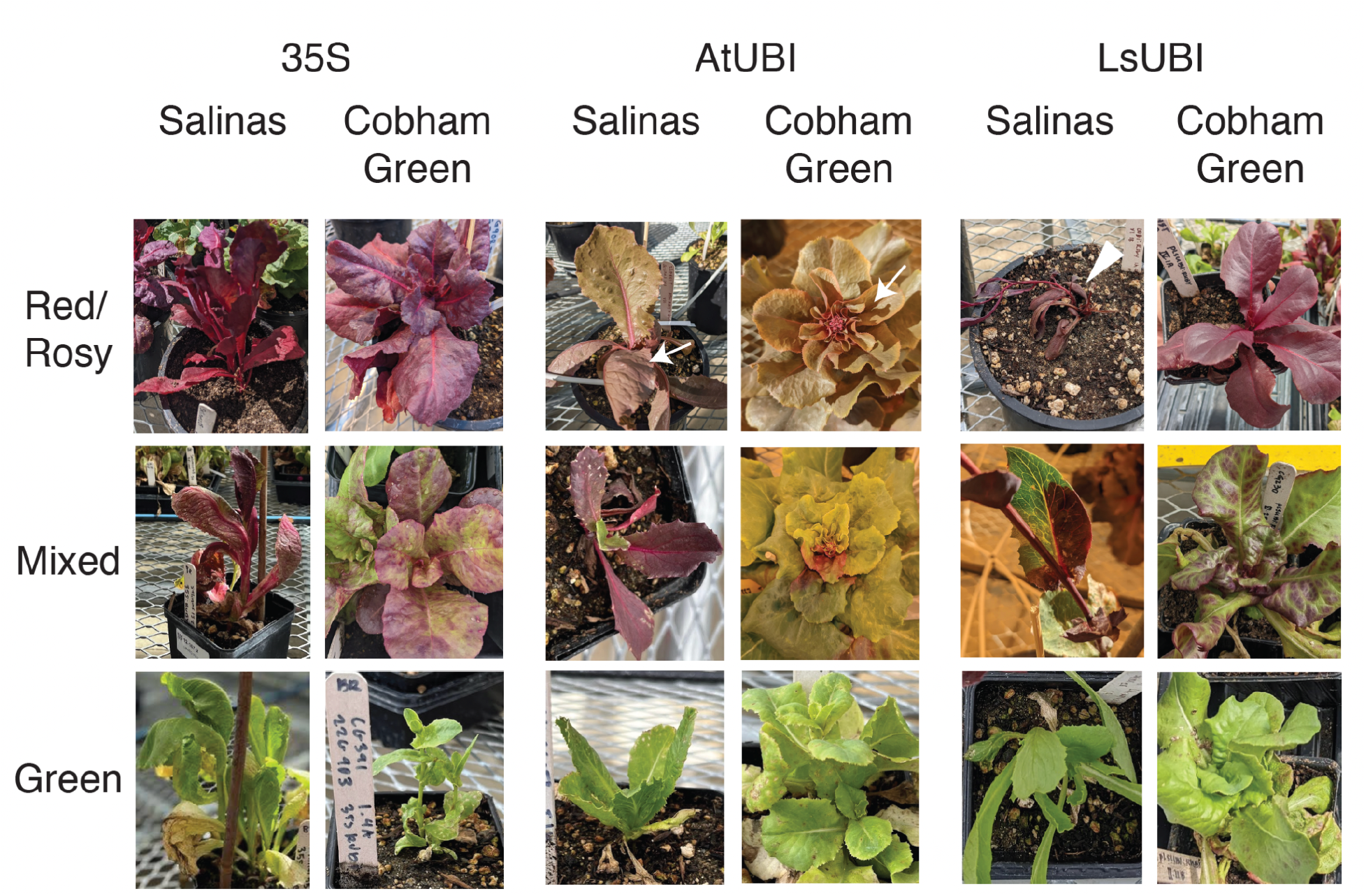
Representative phenotypes of T_1_generation plants from all three RUBY expression constructs in Salinas and Cobham Green. White arrows indicate the lighter “Rosy” phenotype observed in plants transformed with AtUBI::RUBY during vegetative growth in the greenhouse. White arrowhead indicates a representative “Red” LsUBI::RUBY Salinas plant declining after transplanting. This plant and all others in this category did not survive to maturity.

Only about half of the AtUBI::RUBY transgenics with uniform RUBY expression were a deep red color, while the other half of plants with uniform expression exhibited a lighter pink color (Figure 5) that we termed Rosy. Upon transplanting to soil, all Red LsUBI::RUBY Salinas plants failed to survive to maturity (Figure 5). In contrast, six Red LsUBI::RUBY Cobham Green transgenics that were transplanted to soil that survived to maturity. For these transgenics, strong RUBY expression was observed in all parts of the plant throughout vegetative growth and flowering (example shown in Figure 4). Although the LsUBI::RUBY Cobham Green plants with strong RUBY expression grew well, only three of the six independent transgenics produced seeds, which were limited to around 50 or fewer (data not shown). This suggests that RUBY expression imparts a penalty in lettuce that is dependent on both background genotype and developmental stage. We obtained seeds from the three of the sterile or semi-sterile Red LsUBI::RUBY Cobham Green when crossed with wild-type Cobham Green as the female parent, but not as the male parent (data not shown). This suggests that the reason for reduced fertility of plants with high RUBY expression may be explained by pollen sterility, which was surprising given that the pollen did not appear to contain high levels of RUBY (Figure 4C, white arrows).

A subset of plants representing Red, Mixed, and Green phenotypes for all constructs was tested by PCR to assess the proportion of transgenic plants, especially for plants with a Green phenotype (Figure S1). All plants tested positive for the kanamycin selection marker (kan), indicating no “escapes” and that all plants were transgenic. Plants with visible RUBY expression also tested positive for RUBY in all but one case, which was likely a false negative PCR reaction. The majority of Green plants were positive for both RUBY and kan, but there were four cases where plants that were positive for kan were negative for RUBY. Plants testing positive for kan and negative for RUBY may indicate either false negative PCR results or incomplete T-DNA integrations. Previous work in lettuce has shown incomplete T-DNA integrations at similar frequencies (Bull et al., 2023).

### Evaluation of RUBY expression in the progeny of primary transgenics (T_2_ generation)

We obtained seeds from the primary transgenics for all three RUBY expression constructs and for all phenotypic categories. There are three phenotypic color categories each for 35S::RUBY and LsUBI::RUBY (Green, Mixed, Red) and four categories for AtUBI::RUBY (Green, Mixed, Rosy, and Red, where Mixed refers to plants with either mixed Green/Red or mixed Green/Rosy phenotypes), for a total of 10 color/construct categories. T_2_ seeds for both Salinas and Cobham Green were not available for all 10 color/construct categories, either due to a lack of survival or sterility of the T_1_ parents. Since both lettuce cultivars produced similar RUBY expression patterns (Figures 2 and 3), we selected representative transgenic lineages (derived from self-fertilization of 63 T_1_ plants) of either Cobham Green, Salinas, or both for each of the 10 color/construct combinations (Table 2). We sowed 18-21 progeny seeds from each and evaluated the phenotypes at 20 days after sowing. Of a total of 33 T_2_ plants derived from the three semi-sterile Red LsUBI::RUBY T_1_ plants, 24 were Red and 9 were Green, and none were Mixed (Figure 6). This is consistent with the expected 3:1 segregation if the transgenes were inserted at single loci and the RUBY phenotype was stable in all three LsUBI:RUBY T_2_ families (X^2^ test p=1). Analyzing each of the three primary transgenic T_2_ families independently also did not differ significantly from a 3:1 Red:Green distribution; however, the sample numbers were too low for an accurate X^2^ test.

**Figure 6.**
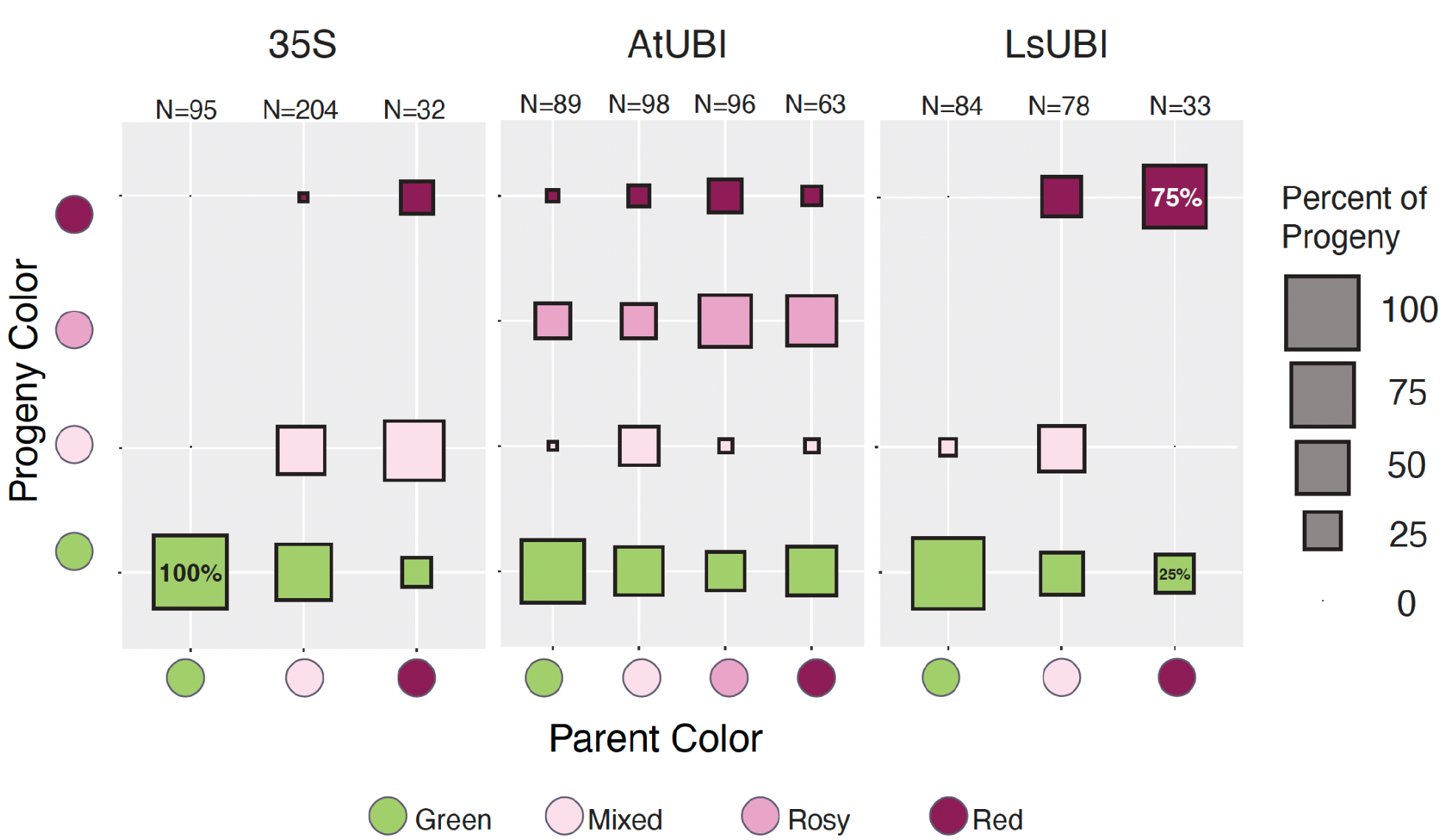
Color phenotypes of progeny by parental phenotype. The percentage of progeny with each phenotype is plotted with respect to the parental phenotype. N = Number of progeny analyzed.

Mixed plants were uncommon among T_2_ families from AtUBI::RUBY Red and Rosy T_1_ parents, while Red plants from the 35S::RUBY lines gave rise to mostly Mixed progeny. Excluding the few Mixed plants in the AtUBI::RUBY T_2_ families, the ratio of Red:Rosy:Green plants among the 93 remaining progeny did not deviate from the 1:2:1 distribution expected for single-locus insertions of the transgene (X^2^ test p=0.74), indicating that the Rosy phenotype is likely due to the hemizygous state.

Slightly more than half of the progeny derived from 35S::RUBY Mixed T_1_ parents were Green and slightly less than half were Mixed. However, individual plants within T_2_ families from AtUBI::RUBY and LsUBI::RUBY Mixed T_1_ parents were distributed more evenly across all phenotypic categories. 35S::RUBY Green T_1_ parents only gave rise to Green progeny, while a small proportion of progeny from Green T_1_ plants from the other two RUBY expression constructs exhibited Red or Mixed phenotypes. This could reflect residual expression of RUBY in a subset of L2 cells in the floral meristem that gave rise to gametes expressing RUBY in the AtUBI::RUBY and LsUBI::RUBY plants that were otherwise phenotypically Green.

We also evaluated 47 progeny from crosses between two independent Red LsUBI::RUBY Cobham Green T_1_ plants and wildtype Cobham Green. Among the 47 progeny scored, approximately 23 were Red and 24 were Green and there were no Mixed plants, consistent with the expectation of a 1:1 segregation of the transgene with no silencing (X^2^ test p=1). Analyzing progeny from each of the two crosses independently also did not statistically significantly differ from the expected 1:1 Red:Green distribution in both cases, but the sample numbers were too low for an accurate X^2^ test for one of the populations because only 11 total progeny seedlings were obtained. The phenotypic distributions are consistent with the transgenes existing as single insertions and we conclude that the RUBY phenotype is stable for LsUBI::RUBY TBC_1_ plants.

### Evaluation of RUBY expression in the T_3_ generation

To determine whether the phenotypes observed in the T_2_ generation were stable in the T_3_ generation, we evaluated the phenotypes of self-progeny of T_2_ plants derived from multiple independent T_1_ plants. For LsUBI::RUBY and AtUBI::RUBY, we selected only Red or Rosy T_2_ parents for phenotypic assessment of plants in T_3_ generation. Since there were only two Red plants in the T_2_ generation for the 35S::RUBY transgenics with available seeds, we also included T_3_ families derived from Mixed T_2_ plants.

Whether the T_3_ families were derived from Red T_2_ plants that had come from Red or Mixed LsUBI::RUBY T_1_ plants, most T_3_ seedlings had a Red phenotype and there were Green seedlings segregating in all T_3_ families (Figure S2). This is consistent with an interpretation that all of the LsUBI::RUBY T_2_ parents were hemizygous for the transgene, but we cannot exclude that the Green plants may have resulted from silencing. The minor proportion of Mixed plants may be taken as an indication that there is minimal ongoing silencing among the T_3_ plants from Red LsUBI::RUBY T_2_ plants, even if the T_1_ progenitor plant had a Mixed phenotype. Thus, we favor the conclusion that Green plants in LsUBI::RUBY T_3_ generation plants reflect null segregants of the T-DNA rather than complete silencing of the RUBY transgene.

For two of the six T_3_ families derived from Red AtUBI::RUBY T_2_ parents, all T_3_ individual plants were Red, indicating that the parents were likely homozygous for the T-DNA. Among the four other T_3_ families derived from Red AtUBI::RUBY T_2_ parents, most T_3_ individual plants were Red, with a minor proportion of Green T_3_ individuals and very few Mixed T_3_ individuals. Of four T_3_ families from Rosy AtUBI::RUBY T_2_ parents, two had mostly or all Rosy plants, while the other two had mostly Green plants. This suggests that Red and Rosy phenotypes of AtUBI::RUBY T_2_ plants largely persisted in the T_3_ generation. In comparison, of the two T_3_ families from Red 35S::RUBY T_2_ plants, one had all Red plants and the other had a broader mix of phenotypes. T_3_ families derived from Mixed 35S::RUBY T_2_ plants commonly had both Green and Mixed plants, suggesting decreasing persistence of RUBY expression in the T_3_ generation for 35S::RUBY plants.

Overall, progeny phenotypes reflected the general patterns observed in the previous generation, with ongoing silencing for lineages that were already experiencing silencing in the T_1_ generation. The majority of 35S::RUBY lineages had moderate to complete silencing by the T_3_ generation (Figure S2 and Table 3). Lineages from Red LsUBI::RUBY T_1_ generation plants showed stable expression in the T_2_, TBC_1_, and T_3_ progeny (Figures 4-7 and S2), with only a small fraction of progeny showing any indication of transgene silencing. AtUBI::RUBY lineages from T_1_ generation Red or Rosy plants generally showed stable expression in the T_2_ generation (Figures 6 and 7), and only one lineage derived from a T_1_ plant with a Rosy phenotype showed reversion to a Green phenotype in the T_3_ generation (Figure S2). We conclude that both LsUBI::RUBY and AtUBI::RUBY exhibited robust RUBY expression in the primary transgenics and over two sexual generations.

**Table 3.** Progeny seeds selected from self-progeny of primary transgenics (T_3_generation)

|  | Background Genotype | T<sub>2</sub> Parent Color<sup>a</sup> | Number of T<sub>2</sub> individuals | Number of T<sub>1</sub> lineages represented<sup>b</sup> | Number of T<sub>3</sub> families selected |
| --- | --- | --- | --- | --- | --- |
| 35S::RUBY | Cobham Green |  |  |  |  |
|  |  | Green | 23 | 5 | 0 |
|  |  | Mixed | 12 | 4 | 2 |
|  |  | Red | 1 | 1 | 0 |
|  | Salinas |  |  |  |  |
|  |  | Green | 23 | 5 | 0 |
|  |  | Mixed | 36 | 7 | 2 |
|  |  | Red | 2 | 2 | 2 |
| AtUBI::RUBY | Cobham Green |  |  |  |  |
|  |  | Green | 16 | 4 | 0 |
|  |  | Mixed | 4 | 3 | 0 |
|  |  | Rosy | 11 | 6 | 2 |
|  |  | Red | 26 | 5 | 5 |
|  | Salinas |  |  |  |  |
|  |  | Green | 16 | 3 | 0 |
|  |  | Mixed | 10 | 3 | 0 |
|  |  | Rosy | 19 | 8 | 2 |
|  |  | Red | 12 | 9 | 1 |
| LsUBI::RUBY | Cobham Green |  |  |  |  |
|  |  | Green | 12 | 4 | 0 |
|  |  | Mixed | 11 | 3 | 0 |
|  |  | Red | 38 | 6 | 9 |
|  | Salinas |  |  |  |  |
|  |  | Green | 6 | 1 | 0 |
|  |  | Mixed | 4 | 1 | 0 |
|  |  | Red | 4 | 1 | 1 |

<sup>a</sup>Color phenotypes of individual T<sub>2</sub> plants were evaluated when plants were entering the seed production phase of their life cycle.
<sup>b</sup> Indicates the number of primary transgenics with T<sub>2</sub> progeny individuals selected for propagation to the T<sub>3</sub> generation.

**Figure 7.**
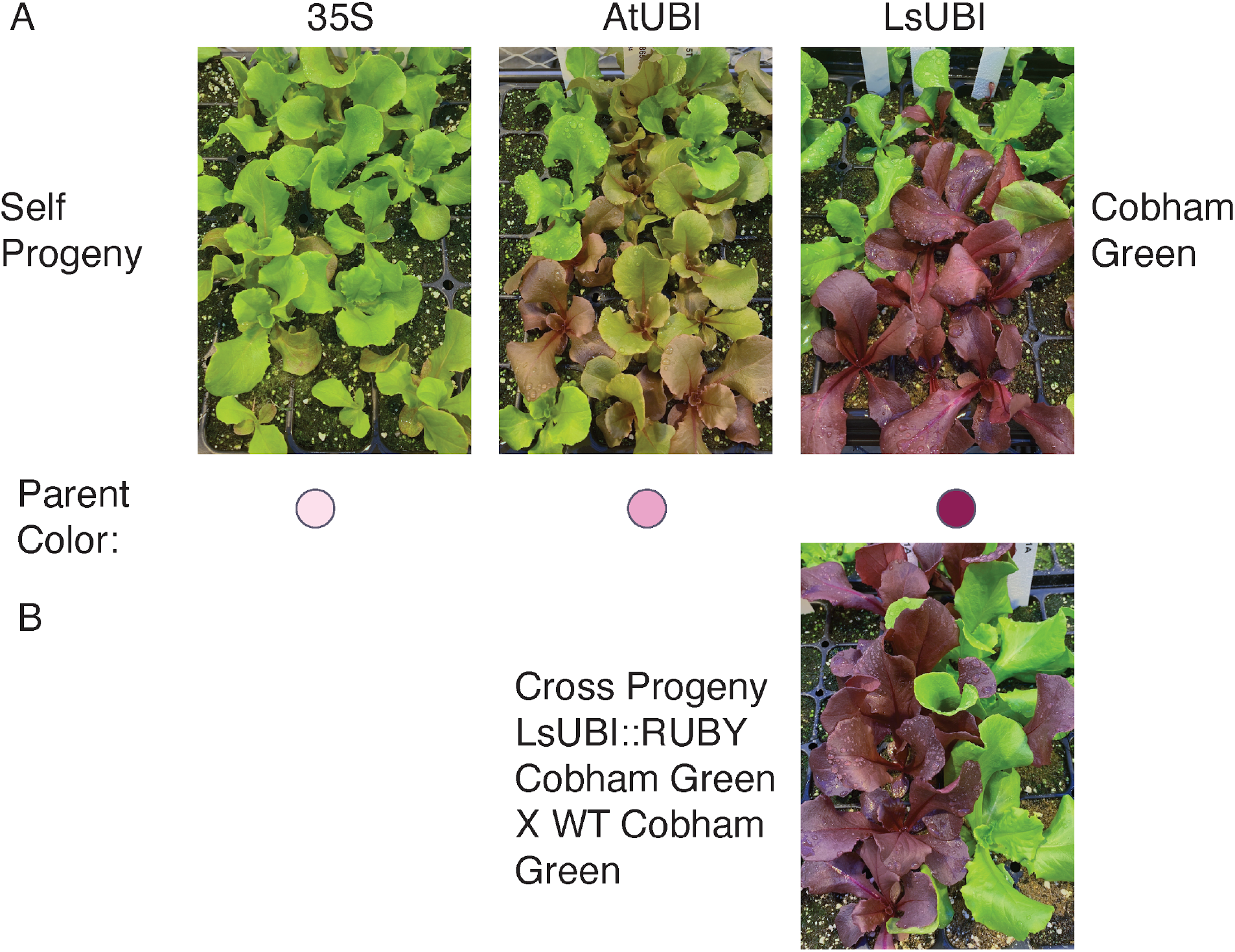
Representative phenotypes of T_2_progeny and progeny from crosses with WT Cobham Green. **A)** T_2_ progeny from self fertilization of T_1_ primary transgenics. Images are of progeny from a single T_1_ primary transgenic for each construct. **B)** Progeny from a cross between LsUBI::RUBY T_1_ plants and WT Cobham Green. Image is of progeny from a single cross that was representative of all progeny from two crosses between these two genotypes.

## Discussion

Here we describe the use of the RUBY reporter to evaluate constructs for achieving the stable expression of transgenes in lettuce. Unlike fluorescent marker proteins, RUBY does not require specific excitation wavelengths or emission filters to be detected and is not subject to photobleaching. This allowed for easy screening of expression phenotypes throughout tissue culture regeneration and at the whole plant level across multiple generations. The LsUBI::RUBY construct showed a strong, uniform expression of RUBY starting at the callus stage that persisted throughout the entire regeneration process and through the next two generations of plants, consistent with Hirai et al. (2011). A lighter phenotype, which we termed Rosy, was only observed for plants transformed with AtUBI::RUBY. The observation of a 1:2:1 segregation of Red:Rosy:Green progeny suggests that the Rosy phenotype could indicate lower overall expression of RUBY from the AtUBI promoter when hemizygous and stronger expression when homozygous due to the increased transgene dosage. Therefore, one potential advantage of the AtUBI promoter is that both strong and intermediate levels of transgene expression could be obtained from the same T-DNA, depending on its dosage.

Plants with Mixed phenotypes were observed for all three constructs in all three generations. These phenotypes could reflect transgene silencing, incomplete integration, tissue chimerism, or the influence of the surrounding chromatin into which the transgene was integrated. Incomplete integration, integration site effects, and tissue chimerism have the same probability of occurrence for all three constructs, since they had the same vector backbone and were introduced using the same strain of *A. tumefaciens*. Therefore, we concluded that differences in the proportion of Mixed phenotypes among the three constructs are largely attributable to the initiation of transgene silencing. The 35S::RUBY construct showed rapid silencing, beginning at even the callus stage, consistent with earlier work demonstrating frequent silencing of the 35S promoter in stable lettuce transgenics (Hirai et al., 2011; Okumura et al., 2016). The expression patterns for each of the three constructs were remarkably similar for both horticultural types of lettuce tested, suggesting that these results may be generalizable across lettuce cultivars. However, since LsUBI shows a more intense RUBY phenotype than AtUBI, it is likely to be the better option if the goal is to achieve transgene expression that is both stable and strong over multiple generations. Recently, Cui et al. (2025) found that the promoter from the peanut ubiquitin 4 gene outperformed the 35S promoter for the expression of both GUS and RUBY marker genes in peanut hairy roots, adding further support for the use of native ubiquitin promoters to achieve high expression of transgenes.

The experiments conducted in this study could not distinguish the individual contributions of the promoters and the terminators, because the three constructs used different combinations of promoters and terminators. Recent work has demonstrated an important role for terminators in determining expression stability and the probability of transgene silencing (de Felippes et al., 2020; Kawazu et al., 2019). The relative strengths of both promoters and terminators may also be different in different species. Kawazu *et al*. (2019) found that the LsUBI promoter drove relatively high levels of expression in the heterologous plant *A. thaliana*, but there was an approximately three-fold higher level of GUS expression when the LsUBI promoter was used with the LsUBI terminator, compared with the NOS terminator. According to de Felippes *et al*. (2020), the *A. thaliana* terminator *RBCS* was associated with a high level of siRNA production and transgene silencing when used in *Nicotiana benthamiana*, but not in *A. thaliana*. Future work evaluating the same promoters with different terminators will further clarify the roles of these two regulatory elements in determining the long-term stability of transgene expression in lettuce.

Despite the advantages of using RUBY compared to other markers, there was a noticeable negative impact on plant growth and fertility in our experiments with lettuce. For Cobham Green, plants expressing RUBY from the LsUBI promoter resulted in low fecundity of the T_1_ transgenics, with all Red plants producing few or no seeds. The same construct in Salinas resulted in a high proportion of deeply red plants, all of which failed to survive transplanting to soil. We observed Red T_2_ plants that were apparently homozygous from the 35S::RUBY and AtUBI::RUBY constructs (based on a lack of segregation in T_3_ progeny), but did not observe any apparently homozygous Red T_2_ plants for LsUBI::RUBY. This may reflect a negative impact on growth or fertility when the LsUBI::RUBY T-DNA is homozygous. The RUBY reporter has been transformed into many different plant species, including *A. thaliana* (He et al., 2020; Yu et al., 2023),cotton (Ge et al., 2023), carrot (Deng et al., 2023), rice (He et al., 2020), poplar (Yuan et al., 2022), tomato (Wang et al., 2023; Yang et al., 2023), maize (Lee et al., 2023; Wang et al., 2023), tobacco (Jogam et al., 2024), *Nicotiana benthamiana* (Yu et al., 2023), *Plukenetia volubilis* (Yu et al., 2023), and the liverwort, *Marchantia polymorpha* (Tse et al., 2024). Most of these studies reported no adverse effects on growth in transgenic plants expressing RUBY. However, in maize, stunted growth was observed in plants expressing RUBY using the 35S promoter (Lee et al., 2023), but not when using the ZmUBI promoter (Wang et al., 2023). Wang et al. (2023) reported negative impacts on growth for 35S::RUBY tomato plants. A study of promoters using the RUBY reporter in the liverwort, *Marchantia polymorpha*, also found that high levels of RUBY expression driven by the 35S promoter had a negative impact on growth, while promoters with intermediate levels of expression and an inducible heat-shock promoter, did not impose the same burden (Tse et al., 2024). Taken together, this suggests that negative impacts of the RUBY marker likely occur when stronger promoters drive a higher level of betalain production and species may be differentially sensitive to this metabolic cost. The RUBY marker is a valuable tool and can be a useful reporter for evaluating how different regulatory elements drive gene expression. However, if the goal is to use RUBY as a transformation marker, researchers should consider using a promoter/terminator combination with an intermediate level of expression or using an inducible system to avoid any potential impacts on growth or fertility.

Our study demonstrated the utility of the RUBY marker for monitoring transgene expression in lettuce. This provided valuable information for selecting promoter/terminator combinations that are unlikely to result in transgene silencing, facilitating future research on engineering traits in lettuce. The RUBY marker will also likely aid the identification of tissue- and cell-specific promoters, or developing environmental sensors by coupling RUBY to the promoters of genes that respond to abiotic and biotic stressors, thus furthering innovations in engineering lettuce.

## Author Contributions

BR and RWM: Experimental design and conception of ideas

BR and CH: Cloning AtUBI and LsUBI RUBY constructs

BR, WR, TB, and CH: Plant transformation

BR, MR, DW, KT, CS, and CH: Phenotypic scoring

BR, DW, MR: Tissue culture

BR: Data analysis and PCR

BR: Wrote manuscript and prepared figures with input from all authors

## Acknowledgements

We thank Guilia Caliandro and Maria J. Truco for assistance with growing plants and making crosses. Juan Debernardi kindly provided us with the 35S::RUBY construct with the kanamycin plant selection marker.

## Material availability

All plasmids associated with this project will be available in Addgene (Table S1).

