## Supplemental Figures S1 and S2 and Supplemental Table S1 for "Monitoring the stability of transgene expression in lettuce using the RUBY reporter"

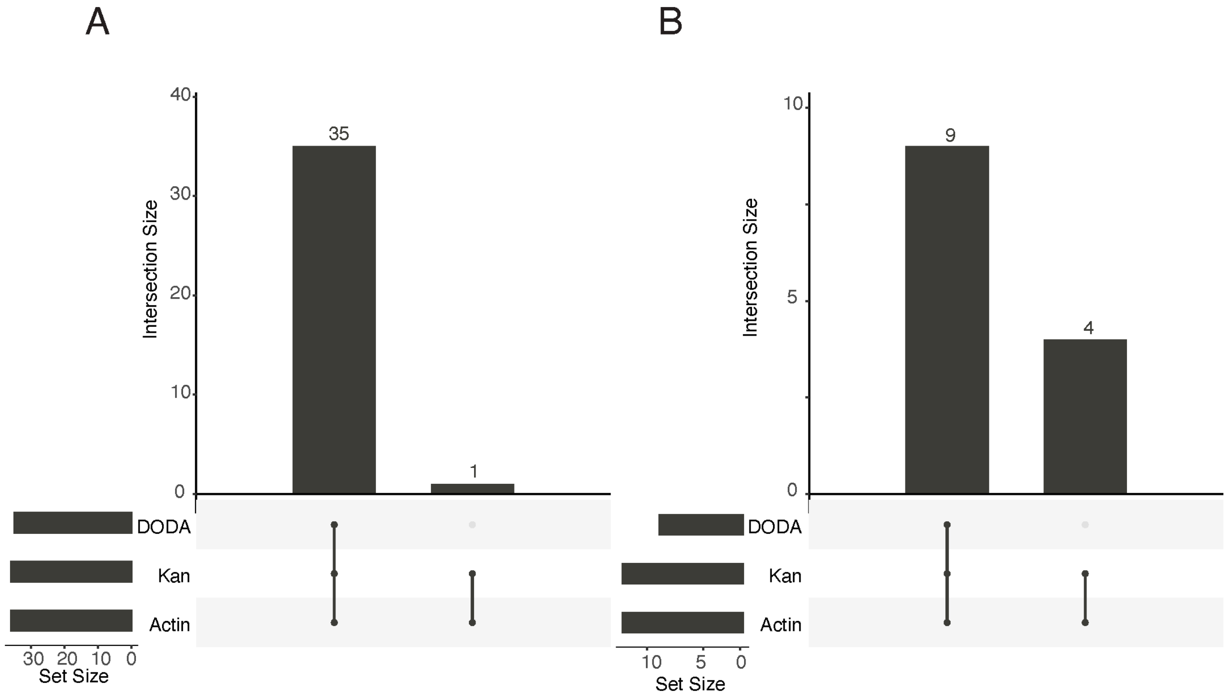


**Figure S1. Analysis of transgene integration in $T_1$ plants by PCR. A)** Combined results across both genotypes and all three RUBY expression constructs for all plants that exhibited at least some RUBY expression. **B)** Combined results across both genotypes and all three RUBY expression constructs for all plants that were entirely green with no RUBY expression. DODA: PCR fragment from the DODA gene of the RUBY expression unit. Kan: PCR fragments for the kanamycin resistance plant selection marker. Actin: PCR fragment for endogenous lettuce control gene.

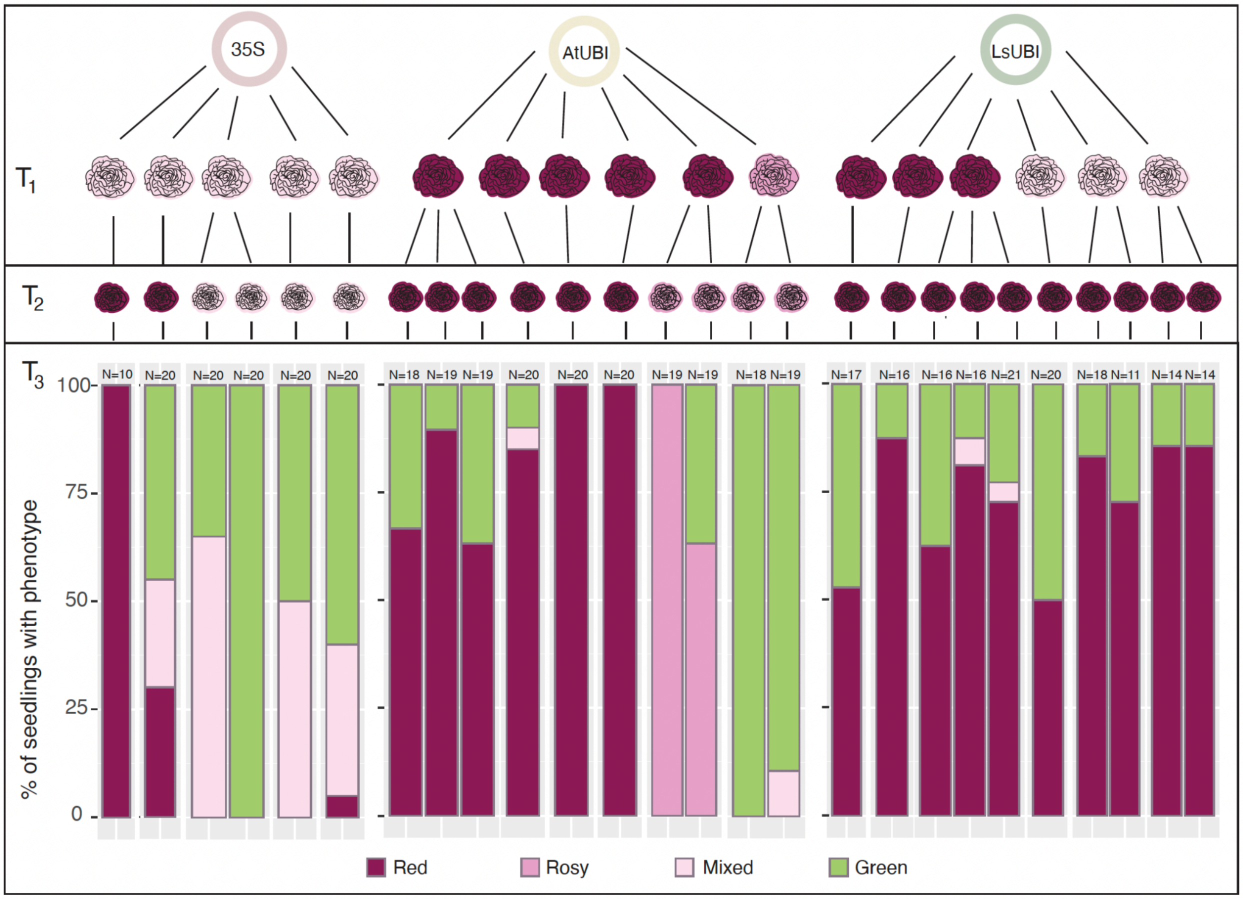


**Figure S2: Phenotypic distribution of $T_3$ progeny.** Seeds of $T_2$ plants were germinated on filter paper in petri dishes and evaluated phenotypically after 7 days. For the AtUBI::RUBY transgenics, we selected $T_2$ plants that had a stable red or rosy phenotype for evaluation in the $T_3$ generation. For the LsUBI::RUBY transgenics, we evaluated the lineages from three mixed and three stable red $T_1$ lines, but only analyzed progeny from stably red $T_2$ plants. Since there were few stably red plants in the $T_2$ generation for the 35S::RUBY transgenics and there was seed available for only one of them, we evaluated one stable red line and four mixed lines. N = number of seedlings analyzed for each $T_3$ family.

**Table S1. Constructs associated with this publication available in Addgene**

| Plasmid Name | Description | Addgene Plasmid ID |
|---|---|---|
| pL0M-PU-pLsUBI | Level 0 GG module with the promoter from the Lactuca sativa polyubiquitin 4 gene. | 244448 |
| pL0M-T-tLsUBI | Level 0 GG module with the terminator from the Lactuca sativa polyubiquitin 4 gene. | 245484 |
| pL0M-SC-RUBY | Level 0 GG module with the RUBY coding sequence. | 245485 |
| pR2B5 | Level 1 transcription unit for pAtUBI-RUBY-tRBCS (R2 position in Golden Gate Level 2 Vector) | 245486 |
| pR2B6 | Level 1 transcription unit for pLsUBI-RUBY-tLsUBI (R2 position in Golden Gate Level 2 Vector) | 245487 |
| pATUBR | Binary plasmid with pAtUBI-RUBY-tRBCS for plant transformation (kanamycin plant selection marker) | 245488 |
| pLSUBR | Binary plasmid with pLsUBI-RUBY-tLsUBI for plant transformation (kanamycin plant selection marker) | 245489 |
| pJD849 | Binary plasmid with p35S-RUBY-tHSP for plant transformation (kanamycin plant selection marker) | 245490 |
